# Dissecting contributions of pulmonary arterial remodeling to right ventricular afterload in pulmonary hypertension

**DOI:** 10.1101/2024.08.18.608471

**Authors:** Sunder Neelakantan, Emilio A. Mendiola, Byron Zambrano, Alexander Vang, Kyle J. Myers, Peng Zhang, Gaurav Choudhary, Reza Avazmohammadi

## Abstract

Pulmonary hypertension (PH) is defined as an elevation in the right ventricle (RV) afterload, characterized by increased hemodynamic pressure in the main pulmonary artery (PA). Elevations in RV afterload increase RV wall stress, resulting in RV remodeling and potentially RV failure. From a biomechanical standpoint, the primary drivers for RV afterload elevations include increases in pulmonary vascular resistance (PVR) in the distal vasculature and decreases in vessel compliance in the proximal PA. However, the individual contributions of the various vascular remodeling events toward the progression of PA pressure elevations and altered vascular hemodynamics remain elusive. In this study, we used a subject-specific one-dimensional (1D) fluid-structure interaction (FSI) model to investigate the alteration of pulmonary hemodynamics in PH and to quantify the contributions of vascular stiffening and increased resistance towards increased main pulmonary artery (MPA) pressure. We used a combination of subject-specific hemodynamic measurements, ex-vivo mechanical testing of arterial tissue specimens, and ex-vivo X-ray micro-tomography imaging to develop the 1D-FSI model and dissect the contribution of PA remodeling events towards alterations in the MPA pressure waveform. Both the amplitude and pulsatility of the MPA pressure waveform were analyzed. Our results indicated that increased distal resistance has the greatest effect on the increase in maximum MPA pressure, while increased stiffness caused significant elevations in the characteristic impedance. The method presented in this study will serve as an essential step toward understanding the complex interplay between PA remodeling events that leads to the most severe adverse effect on RV dysfunction.

## 1 Introduction

Pulmonary hypertension (PH) is characterized by an elevation in mean pulmonary arterial pressure (mPAP) and affects *∼*1% of the world population^1^. PH can be subdivided into several etiologies. In the case of group 1 PH, known as pulmonary arterial hypertension (PAH), the increase in mPAP, and subsequently, the right ventricular (RV) afterload, is due to an increase in pulmonary vascular resistance (PVR) and a decrease in pulmonary arterial compliance (PAC)^2, 3^. These changes are primarily linked to the decreased compliance of the proximal arteries and increased resistance in the distal pulmonary vasculature, caused by the narrowing of the vessels and the “pruning” of distal arteries^1, 4^. In turn, the elevations in RV afterload lead to increased RV wall stress^5, 6^, triggering a cascade of mechano-driven RV remodeling events^7–11^.

If left untreated, RV remodeling can lead to right heart failure and death. The standardized death rate in PH was reported to be 4.5 to 12.3 in a 100,000 population^12^. The median survival was reported to be 7 years for patients with PAH and 2.8 years for patients with idiopathic PAH^13–15^. As RV remodeling is primarily triggered by increases in RV afterload, understanding the individual contribution of each vascular remodeling event toward the elevations of RV afterload is essential. Also, as each pulmonary arterial (PA) remodeling event may require different clinical intervention strategies, there is a need to develop tools that can quantify the effects of isolated remodeling events on RV afterload and PA pressure in a patient-specific manner to enable optimized treatments. The development of such tools remains an unmet need in PH.

The primary tissue-level remodeling events in the pulmonary artery (PA) driving increased PA pressure and RV afterload are increased resistance caused by reduced lumen area in the distal vessels and decreased vessel compliance (increased vessel stiffness). Increased resistance is predominantly caused by the narrowing of distal vessels, including precapillary arterioles^16, 17^. Increased vessel stiffness occurs through smooth muscle hypertrophy and proliferation of vascular cells, which contribute to increased deposition of extracellular matrix components. However, distinguishing the contributions of resistance and compliance to afterload remains difficult due to the inadequacy of existing in-vivo indices that aim to assess these contributions. In contrast, in-vivo measurements, such as catheterization and medical imaging, combined with in-silico modeling^18–20^, offer an innovative approach to investigate patient-specific pulmonary hemodynamics and quantify the contribution of increased stiffness and resistance.

Several studies^21–26^ have significantly advanced the use of in-silico modeling approach to investigate pulmonary hemodynamics in the past decade. Such studies have developed three-dimensional (3D) computational fluid-structure interaction (FSI) models of the proximal vasculature to analyze hemodynamic behavior in various vascular diseases, including PH. In such studies, the structure of the proximal vasculature is obtained through medical imaging, and the distal vasculature is represented by Windkessel elements. The flow boundary conditions used in such studies are generally obtained through catheterization and/or 4D flow magnetic resonance imaging (MRI). These models^21–26^allow for high-fidelity simulations and the incorporation of subject-specific vascular geometry, enabling the accurate assessment of parameters such as wall shear stress (WSS) and vorticity. Due to their higher fidelity, these models also tend to be better suited to analyze turbulent flow behavior. While such models enable the in-silico simulation of physiologically realistic flow in the vessels, they can be computationally expensive when incorporating the actual geometry of the distal vasculature. This limits the potential application of high-fidelity 3D FSI models in clinical settings. In contrast, other studies have developed reduced-order models to reduce computational time and cost while including the distal vasculature in the models. One such type of reduced order that has recently gained interest, especially in terms of subject-specific simulations, is 1D FSI models^27–33^. Such models tend to approximate vessels as a line with a given length and lumen area. Also, these models work under the assumption of laminar and axisymmetric flow. Studies have demonstrated the ability of 1D FSI models of the vascular system to capture physiologically realistic pressures in the vessels^27, 30^. Such studies have also reported the capacity to include a significant portion of the pulmonary vasculature (up to 400 vessels), with the number of vessels being limited by the imaging modality. Such models with reduced fidelity significantly reduce computational costs while offering an accurate estimation of pressure and vessel stress, facilitating the translation of such models to be translated into clinical applications. However, reduced-order models have yet to be used to address the need for an improved understanding of the contributions of individual PA remodeling events toward altered vascular hemodynamics in PH.

In this study, a recently developed 1D FSI model^27, 30, 31^ was extended to account for PA vessel anisotropic and nonlinear biomechanical behavior, and the extended model was used to separate the effects of alterations in PA vessel biomechanics on pulmonary hemodynamics in a rodent model of PH. The in-silico models were made to be fully subject-specific by integrating the vascular morphology reconstructed using X-ray micro-tomography^34^ of the rat lungs (Fig. 1), the mechanical behavior of the arterial vessels, estimated through the ex-vivo testing of the main PA (MPA), and hemodynamic data obtained using right heart catheterization and Doppler imaging. The in-silico models were able to separate the effect of the following events in MPA pressure elevations and altered pulsatility (quantified by PA impedance): (i) increased proximal stiffness and distal stiffness, and (ii) increased distal resistance. This study is expected to assist with developing image-based strategies to individualize and optimize PA remodeling-targeted treatments based on predicting the effects of individual PA remodeling restoration on MPA pressure reductions in each patient.

**Fig. 1.**
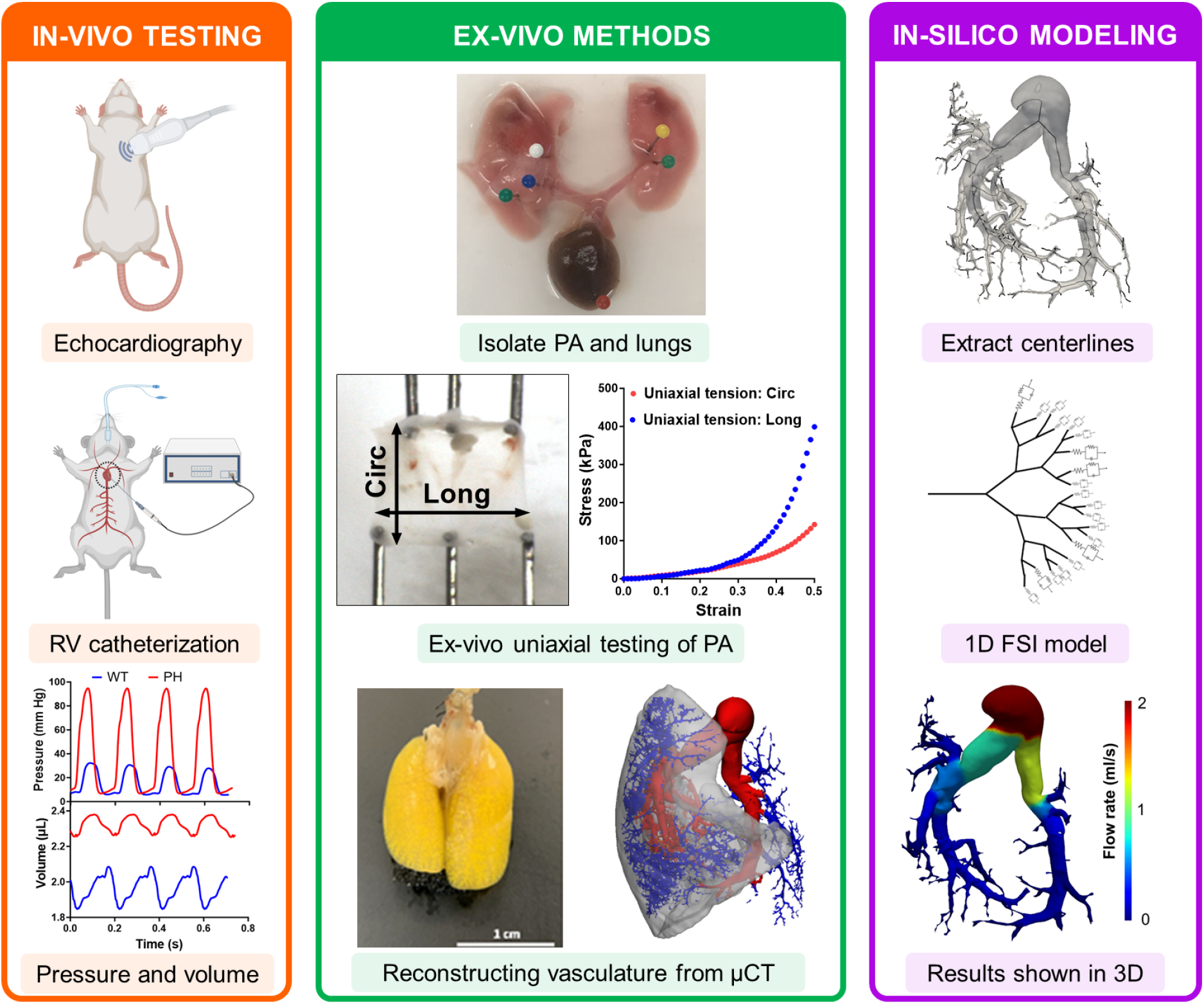
Combined in-vivo, ex-vivo, and in-silico approaches were used to separate the contribution of individual remodeling events towards increased MPA pressure. Data from in-vivo and ex-vivo measurements were combined to create an in-silico model of the pulmonary vasculature. CTL: control, PH: pulmonary hypertension, PA: pulmonary artery, MPA: main pulmonary artery, Circ: circumferential, Long: longitudinal, *µ*CT: X-ray microtomography, FSI: fluid structure interaction.

## 2 Methods

All housing and procedures were performed at the AAALAC-certified and accredited rodent care facility at the Providence VA Medical Center and were performed according to protocols approved by the Providence VA Medical Center’s IACUC (IACUC 1633548).

### 2.1 Animal model of PH

A total of 14 Fischer (CDF) rats were used for this study (n=7 control, n=7 PH). PH was induced via an established Sugen-hypoxia (SuHx) protocol as described by Vang et al.^35^. Briefly, PH was developed by a single subcutaneous injection of a vascular endothelial growth factor inhibitor (SU5416, 20 mg/kg body weight; APExBIO, Houston, TX) that was dissolved in a diluent (0.5% carboxymethylcellulose, 0.9% NaCl, 0.4% Polysorbate 80, and 0.9% benzyl alcohol) followed by three weeks of normobaric hypoxia exposure (10% FIO_2_; A-Chamber Animal Cage Enclosure with ProOx 360 High Infusion Rate O_2_ Controller, BioSpherix Ltd, Parish, NY) and subsequent housing in normoxic conditions for one additional week. The control (CTL) rats received a diluent injection and were housed in normoxic conditions until the end of the study. At the end of the study, animals were placed on a heating pad (37°C) and anesthetized with continuous isoflurane inhalation (1.5%–2%) in 100% O2 for the duration of the echocardiography and catheterization procedures. Transthoracic echocardiography and right heart catheterization were performed as previously described^35^. After catheterization, the rats were euthanized under isoflurane anesthesia by exsanguination. The lungs of one rat from each group (n=1 CTL, n=1 PH) were isolated and used for vascular imaging, described in the following section.

### 2.2 Vascular imaging

The geometry of the pulmonary vasculature was obtained through the process described by Knutsen et al.^34^. Briefly, the rats were euthanized through exsanguination and placed in the supine position with the lungs and trachea exposed. The ribcage of the rat was removed to expose the heart and lungs. A 30G needle was inserted into the right ventricle through the apex, and 10^-4^ M sodium nitroprusside (SNP) in PBS was pumped in to flush the blood and dilate the pulmonary vasculature. Then, the trachea was incised, and formalin was injected using a pressure head of 20cm to ensure that the lungs were fixed in a pressurized and inflated state. The lungs were then excised and isolated. To cast the vasculature, a solution containing 8:1:1 of polymer: diluent: curing agent was mixed (Details of the polymer compound are available in^34^) and infused into the vasculature through the catheter previously inserted into the apex. The compound was allowed to harden for 30-40 min at room temperature, after which the lungs were placed in 10% formalin overnight. For imaging, paraffin film was used to create a flat surface on the scanning bed, and the lungs were imaged using micro-CT to obtain the vascular geometry of the lungs. The images were analyzed using Slicer and the vascular modeling toolkit (VMTK) to obtain the length and radius of the vessels of the pulmonary vasculature (Fig. 2). The VMTK toolkit was used to estimate the end points of individual vessels of the vascular tree. The endpoints of the individual vessels were used to obtain the vessel connectivity information, which, in combination with the vessel dimensions, was used to describe the 1D PA vasculature.

**Fig. 2.**
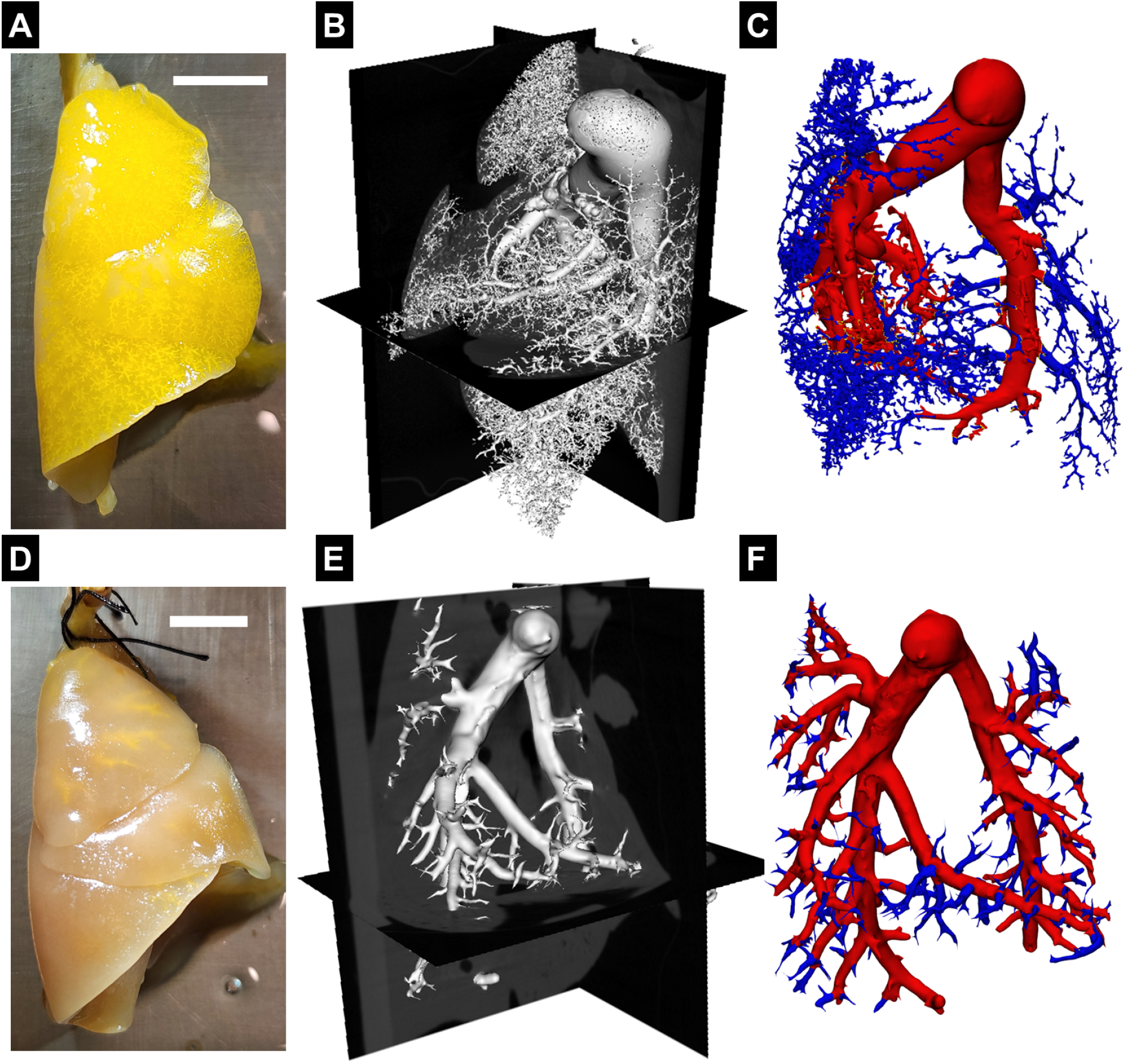
Resin filled lungs from **(A)** control (CTL) rat and **(D)** pulmonary hypertension (PH) rat, Scale bar - 1cm. Representative X-ray microtomography image slices of fixed lungs, with polymer resin infused pulmonary vasculature of **(B)** CTL and **(E)** PH rat. Reconstructed pulmonary vasculature of **(C)** CTL and **(F)** PH rats.

### 2.3 Mechanical testing of PA vessels

The MPAs of the rats, not used for vascular reconstruction (n=6 CTL, n=6 PH), were isolated and used to perform ex-vivo uniaxial mechanical testing. The MPA was cut and opened along its axis to create a rectangular tissue specimen. These specimens were subjected to uniaxial mechanical testing along both directions (circumferential and axial directions of the intact vessel) using a mechanical testing machine (Cellscale Biotester; Fig. 1). The specimens were stretched to a strain of 50% in both directions, and the stress measured from this test has been reported as 1st-Piola Kirchoff (1st-PK) stress. Stiffness was estimated as the slope of the stress-strain curve at 50% strain.

### 2.4 1D FSI model

The flow behavior in the pulmonary vasculature was simulated using a 1D FSI model adapted from Olufsen et al.^27, 30, 31^. This model describes the flow and blood vessel interaction in the pulmonary vasculature (proximal and distal). The 1D vascular tree was connected to a 3-element Windkessel model describing the flow condition in the smaller distal vessels and the capillary beds. This approach assumes that the flow is incompressible, Newtonian, laminar, and axisymmetric. Briefly, the continuity and momentum equations are integrated over the vessel’s cross-sectional area, yielding the following equations

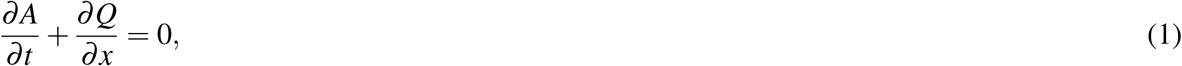

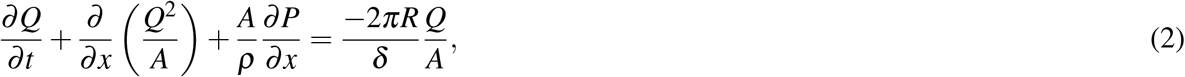

where *x* and *t* denote the spatial and temporal coordinates, *Q*(*x, t*) is the volume flow rate, *P*(*x*,*t*) is the pressure, *R*(*x, t*) is the radius of the vessel and *A*(*x, t*) is the cross-sectional area. 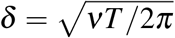 is the boundary layer thickness, where *T* is the time per cardiac cycle. The fluid density *ρ* and kinematic viscosity *ν* are assumed to be constant. The bifurcation of flow was based on satisfying the following conditions

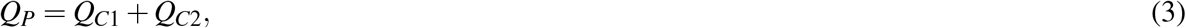

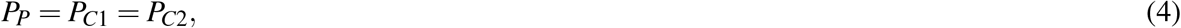

where *Q*_*P*_, *P*_*P*_ are the volume flow rate and pressure at the distal end of the parent vessel, *Q*_*C*1_, *P*_*C*1_, *Q*_*C*2_, *P*_*C*2_ are the volume flow rates and pressures at the proximal end of the branching child vessels. The 3-element Windkessel boundary condition at the terminal vessels to characterize the flow in the capillary beds is given by an RCR circuit model

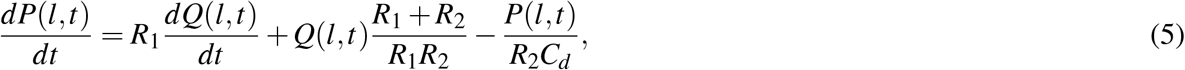

where *R*_1_ and *R*_2_ are resistance elements and *C*_*d*_ is the compliance element. *R*_1_ and *R*_2_ were set to be equal. *R*_1_,*R*_2_,*C*_*d*_ represent the net resistance and compliance of the smaller vessels downstream of the endpoints of the vasculature. Distal stiffness (*S*_*d*_) was defined as the inverse of distal compliance *S*_*d*_ = 1*/C*_*d*_.

The 1D model was extended to account for the vessel tissue’s anisotropic and nonlinear constitutive behavior. A constitutive model of the artery wall was defined to relate the internal pressure with the lumen area. The Holzapfel-Ogden (H-O) model^36^ was used to capture both non-linearity and anisotropy of the passive mechanical behavior of the pulmonary vasculature observed in the ex-vivo experiments. The stress state of the vessel under loading is given by *σ*. Here, 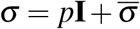, where 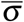 is the isochoric part of the stress tensor and *p* is the hydrostatic pressure. Equations for the isochoric stress terms in the polar co-ordinates are given by

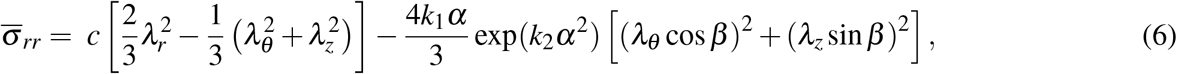

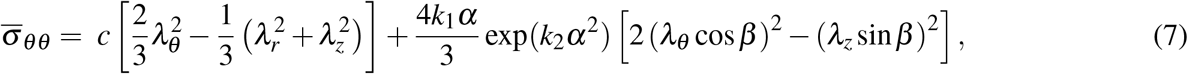

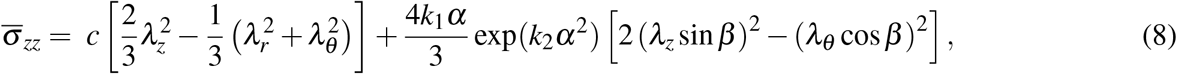

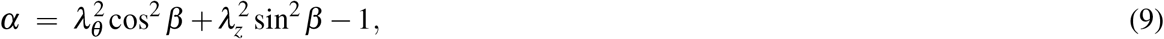

where *λ*_*r*_, *λ*_*θ*_, *λ*_*z*_ are the stretch values in the *r, θ, z* directions respectively. *c, k*_1_, *k*_2_ are the material parameters (detailed in section S.1) that were estimated using the stress-strain data from ex-vivo mechanical testing of the MPA. *β* is the orientation of the muscle and collagen fibers with respect to the circumferential direction. The equilibrium equation in the radial direction was used to determine the fluid pressure,

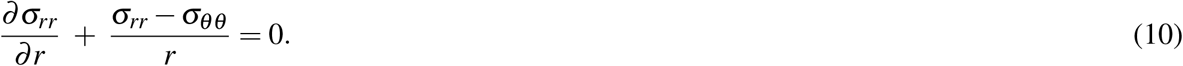

Integrating this equation over the thickness of the vessel, assuming no pressure outside the vessel, yields the pressure in the interior of the vessel, given by

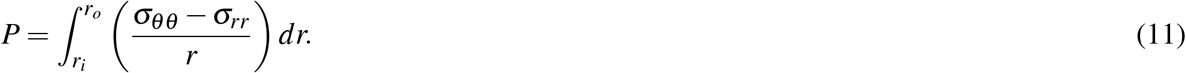

Next, the stress terms are written in terms of the isochoric components as

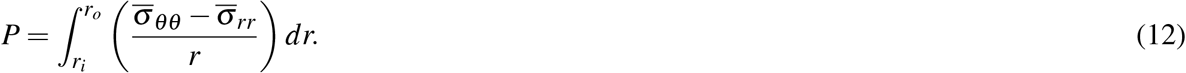

Substituting Eqs. S.25 and S.26,

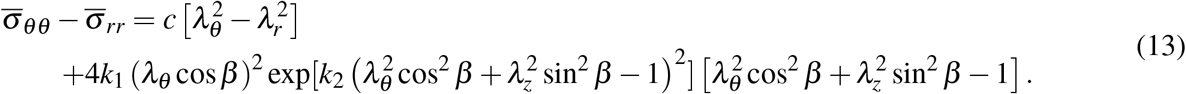

The detailed derivation for Eqs. 6,7,8,13 can be found in section S.1 of the supplementary material. Non-linear regression of the ex-vivo uniaxial testing results was performed to obtain the material parameters introduced in this subsection. When the specimen was stretched along the axial direction, 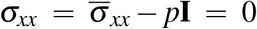 (Eq. 6) and 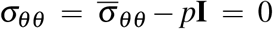 (Eq. 7) conditions were imposed to determine *p* and *λ*_*θ*_ in terms of the material parameters and estimate *σ*_*zz*_ (Eq. 8). This process was repeated for the uniaxial test along the circumferential direction. 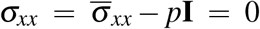 (Eq. 6) and 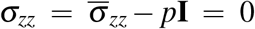 (Eq. 8) conditions were imposed to determine *p* and *λ*_*z*_ in terms of the material parameters and estimate *σ*_*θθ*_ (Eq. 7). The estimated *σ*_*θθ*_ and *σ*_*zz*_ were compared against the measured values to determine the error, which was minimized using least-squares curve fitting.

### 2.5 In-silico experiments

#### 2.5.1 Parameterization of the model

To modulate proximal vessel stiffness, a scaling factor, termed proximal stiffness (*S*_*p*_), was defined that uniformly scales material parameters *c* and *k*_1_ such that *S*_*p*_ = 1 represented mechanical behavior from control specimens (Table. 1). Proximal vessel compliance is inversely proportional to *S*_*p*_. The 3-element Windkessel model represents the flow in the smaller distal vessels and capillary beds. In the Windkessel elements, *R*_*d*_ = *R*_1_ = *R*_2_ represented the hydraulic resistance (flow resistance), and *S*_*d*_ was taken to represent the stiffness of the elastic (distal) arterioles (Table. 1). The effects of these individual remodeling mechanisms were explored, as described in the next section.

**Table 1.** Modeling parameters represent remodeling mechanisms.

| Mechanism | Model Parameter |
| --- | --- |
| Proximal stiffness | $S_p$ |
| Distal stiffness | $S_d$ |
| Distal resistance | $R_d = R_1 = R_2$ |

#### 2.5.2 Deconvoluting the contributions of vascular remodeling events to increased RV pulse pressure

In-silico experiments were performed to separate the contributions of individual vascular remodeling events on the MPA pressure waveform. To begin the in-silico simulation of the CTL rat, the PA constitutive material constants (*c, k*_1_, and *k*_2_) were obtained by fitting the model to the ex-vivo mechanical testing data. The values obtained for each specimen were averaged, and the mean values were used in the in-silico simulations. Next, the distal stiffness (*S*_*d*_) and resistance (*R*_*d*_) were chosen such that the simulated pressure matched the measured pulse pressure at MPA (ΔMPA pressure), defined as the difference between the systolic and diastolic pressures. The rate of change in RV volume was used as the flow rate at the MPA inlet, assuming no regurgitation at the tricuspid valve. The waveform was uniformly scaled such that the area under the curve (inflow volume per pulse) matched the cardiac output measured through echocardiography. In addition, the initial pressure in the simulation was set to zero.

Next, to estimate the alteration in resistance and compliance for the in-silico PH rat, the distal resistance (*R*_*d*_) was scaled based on the change observed in PVR from healthy to PH rats. Distal stiffness (*S*_*d*_) and proximal stiffness (*c*) were scaled based on the increase in ex-vivo mechanical stiffness. The variable ↑ *S* was used to represent the combined increase of both proximal (*c*) and distal (*S*_*d*_) stiffness. After scaling all relevant parameters, the PH model was used to estimate pressure to obtain an in-silico prediction of ΔMPA pressure in PH using the flow waveform scaled by the average cardiac output measured in the PH animals.

#### 2.5.3 Deconvoluting the contributions of vascular remodeling events towards altered pulsatility of flow

While peak pressure is an important metric for understanding the increase in RV afterload, it is insufficient to describe the pulsatile and dynamic behavior of pressure and flow in the pulmonary vasculature and their alterations in PH. The alteration in the pulsatility of the pressure and flow waveforms is commonly analyzed using pulmonary arterial impedance (PAZ) in frequency domain^37–39^. The flow and pressure waveforms were converted to the frequency domain using Fourier transforms, and PAZ was calculated using the following equation,

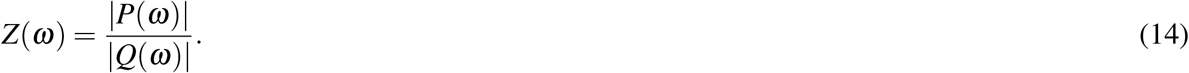

Two major indices of the PAZ include the impedance at 0 Hz, denoted by *Z*_0_, and the characteristic impedance, devoted by *Z*_*c*_ and defined as the average impedance over a large frequency range excluding 0Hz (Fig. S3 shows a representative example). Here, *Z*_*c*_ was calculated over the range of 0-250 Hz. *Z*_0_ and *Z*_*c*_ were used to understand the effect of vascular remodeling on the mean and pulsatile (oscillatory) behavior of the pressure and flow waveforms. *Z*_0_ and *Z*_*c*_ were estimated for four scenarios -control, increased vessel stiffness, increased distance resistance, and PH. Since our key objective here was to separate the effects of increased PA stiffness versus increased PA resistance on the flow pulsatility, the pulse pressure waveform was used in Eq. 14 to estimate impedance. By definition, the inclusion of a diastolic pressure baseline in the pressure waveform would leave *Z*_*c*_ unaltered.

### 2.6 Statistical analysis

The experimental data was analyzed in GraphPad Prism 9. All the measured data were presented as mean *±* standard error. Data from male and female rats included in the study were denoted by filled and hollow markers, respectively. Paired student t-tests were performed when analyzing the statistical significance between the CTL and PH rats in Fig. 4. The numerical significance values have been reported in this study^40^.

## 3 Results

### 3.1 Morphological remodeling of PA vasculature

There was a decrease in the number of smaller vessels in the PH lungs after being filled with resin (Figs. 2B, E). As the resin filling was performed under the same pressure and for the same amount of time, this difference indicated increased resistance in the distal vessels in PH (Figs. 2C, F). Catheterization indicated increased diastolic and systolic pressure (Figs. 3A, B). The increased distal resistance observed during resin filling was reflected in the PVR measured through catheterization (Fig. 3C). Stroke volume decreased (Fig. 3D), which was in agreement with literature^10^. In addition to changes in the geometry, an increase was observed in the MPA thickness (Fig. 3E).

**Fig. 3.**
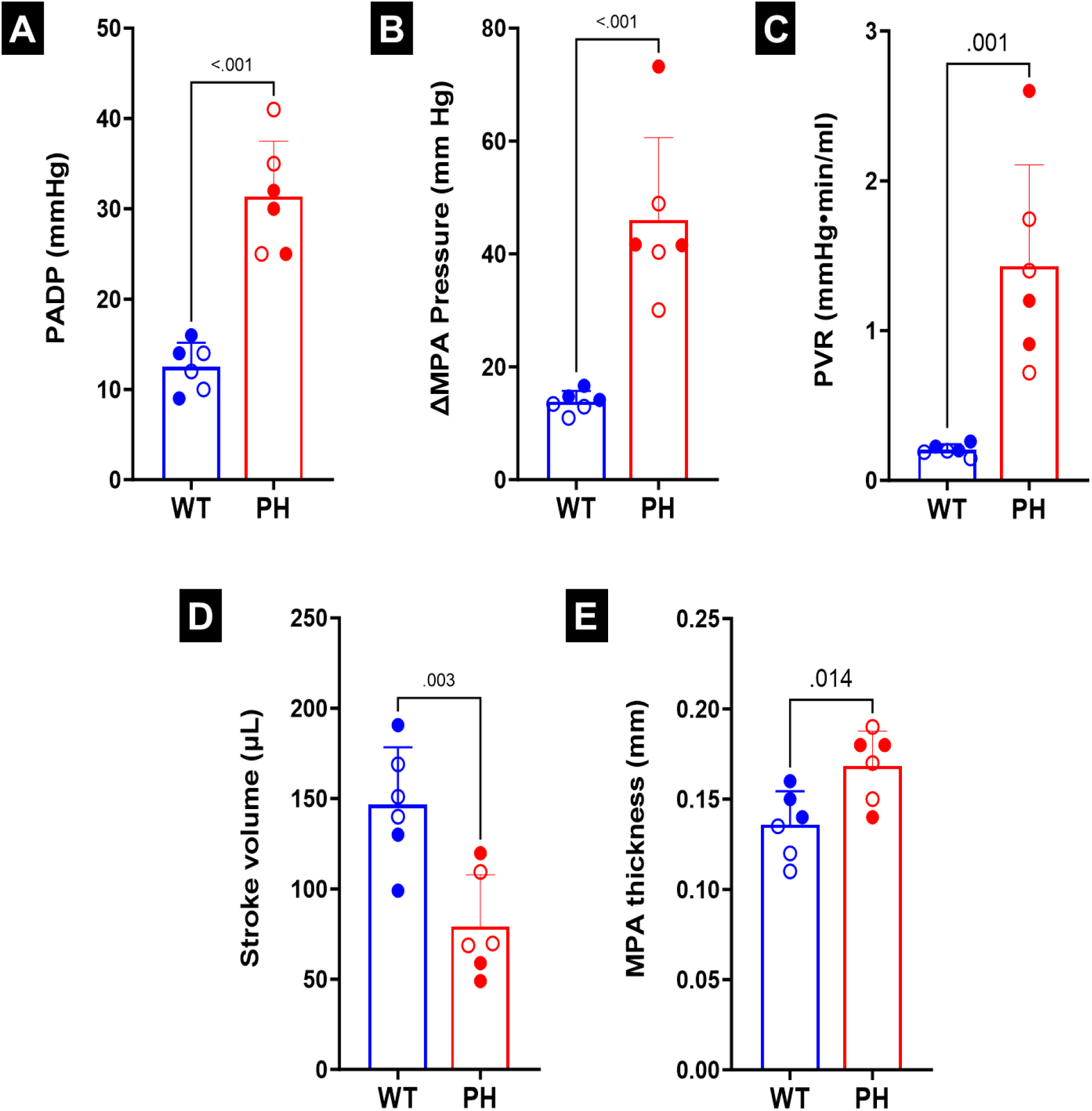
**(A)** Pressure in the main pulmonary artery (MPA) at diastole (PADP). **(B)** Difference in MPA pressure between systole and diastole (ΔMPA pressure).**(C)** Pulmonary vascular resistance (PVR.) **(D)** Stroke volume. **(E)** MPA thickness. Statistics were performed using student t-tests. Male and female animals are denoted by filled and hollow markers, respectively. CTL: n=6, PH: n=6.

**Fig. 4.**
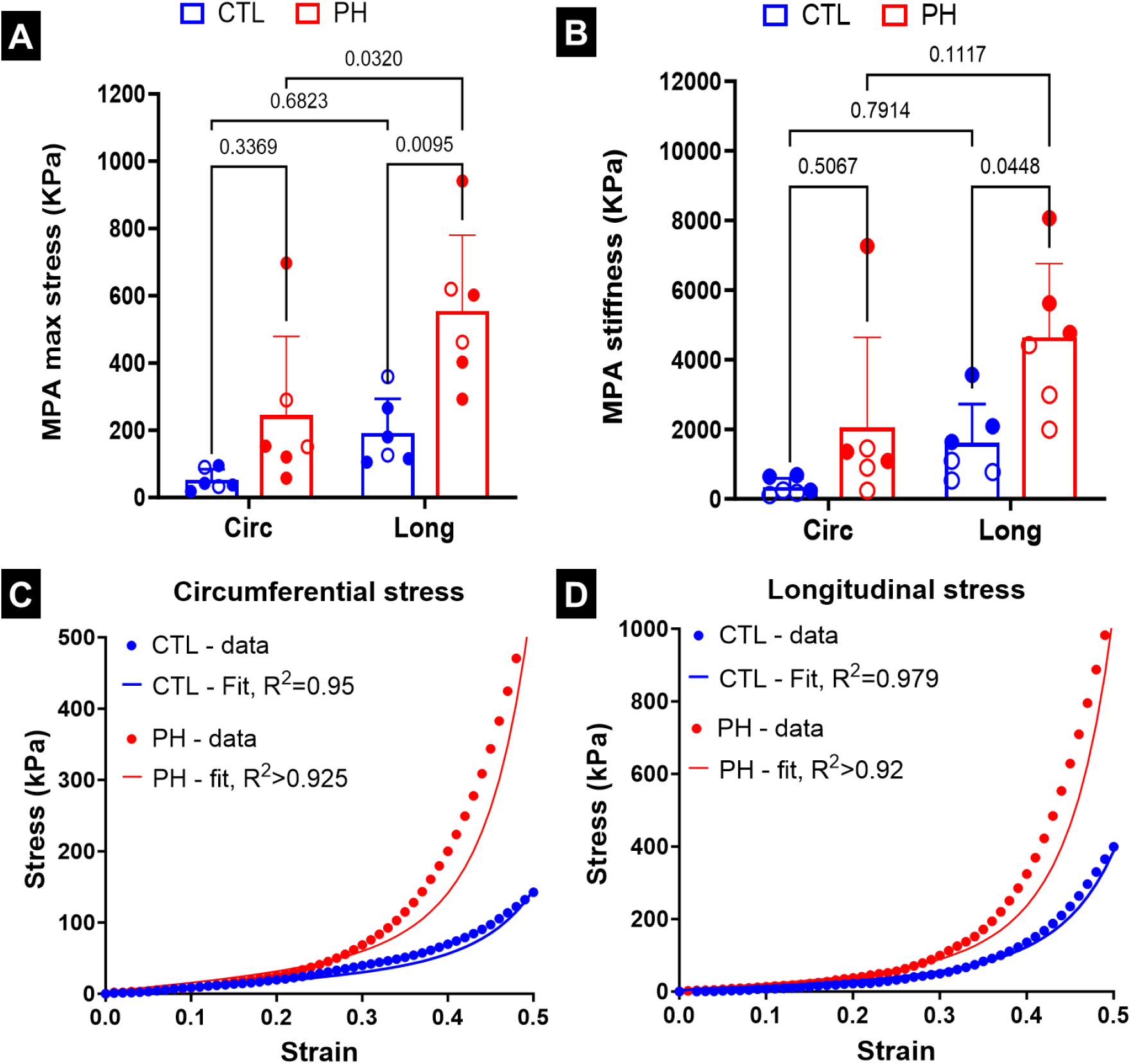
**(A)** Stress in the main pulmonary artery (MPA) during mechanical testing at 50% strain, **(B)** stiffness of the MPA at 50% strain. Mean stress-strain curve based on uniaxial testing of pulmonary arteries and fits based on non-linear artery model in **(C)** circumferential and **(D)** longitudinal directions. Statistics in **(A), (B)** were performed using 2-way ANOVA with Tukey’s correction for multiple comparisons. Male and female animals are denoted by filled and hollow markers, respectively. CTL: n=6, PH: n=6. Circ: circumferential, Long: longitudinal.

### 3.2 Mechanical stiffening of PA tissues in PH

There was a significant increase in the maximum stress (Fig. 4A) and stiffness of the MPA (Fig. 4B) in the longitudinal direction in the PH specimens as compared to the controls. The MPA tissue specimens of the CTL rats were biased towards the longitudinal direction (Fig. S1), but the difference was not significant. The MPA specimens of the PH rats were further biased towards the longitudinal direction (Fig. S1), with this change being significant when comparing the maximum stress (Fig. 4A), and near-significant when comparing the stiffness (Fig. 4B). The non-linear constitutive model proposed in this study was able to closely capture the behavior of the PA specimens observed during ex-vivo uniaxial testing (Figs. 4C, D).

### 3.3 Relationship between vascular remodeling and pulmonary hemodynamics

The measured volume flow rates (Fig. 5A) were used as inputs to investigate the cumulative effects of the PA remodeling events on ΔMPA pressure. There were alterations observed in both the maximum value and relaxation behavior of the pressure waveform (Fig. 5B). When the model resistance, and stiffness were individually modified to match the values observed in the PH rats, the increase in distal resistance had the largest effect towards increased ΔMPA pressure (Fig. 5C). The vessel thickening (observed in Fig. 3E) was incorporated into the increased PA stiffness. Increased vessel stiffness led to smaller and relatively equal increases in ΔMPA pressure (Fig. 5C). In addition, increased stiffness did not cause significant changes to the qualitative behavior of the pressure waveform (Fig. S2). However, increased resistance caused a delay between the peak of the volume flow rate and the pressure waveform and resulted in a significantly slower drop in pressure from the peak (Fig. S2). Next, the cumulative effects of the remodeling events were calculated (Fig. 5D). Expectedly, incorporating decreased flow rate (CTL - PH Flow; Fig. 5D) led to a decrease in ΔMPA pressure. Increased stiffness (*Δ*S; Fig. 5D) led to small increases in ΔMPA pressure, while adding the effect of increased resistance caused a significant increase in ΔMPA pressure (↑S+↑R_d_; Fig. 5D). However, comparing the individual and cumulative effects of the PA remodeling events indicated that a notable portion of the increase in ΔMPA pressure originated from the interactions between these two events (Fig. S2). When the resistance, stiffness, and volume flow rate all matched the corresponding values in PH rats, the ΔMPA pressure value became comparable to the experimentally observed ΔMPA pressure values in the PH rats (↑S+↑R_d_ PH Flow; Fig. 5D).

**Fig. 5.**
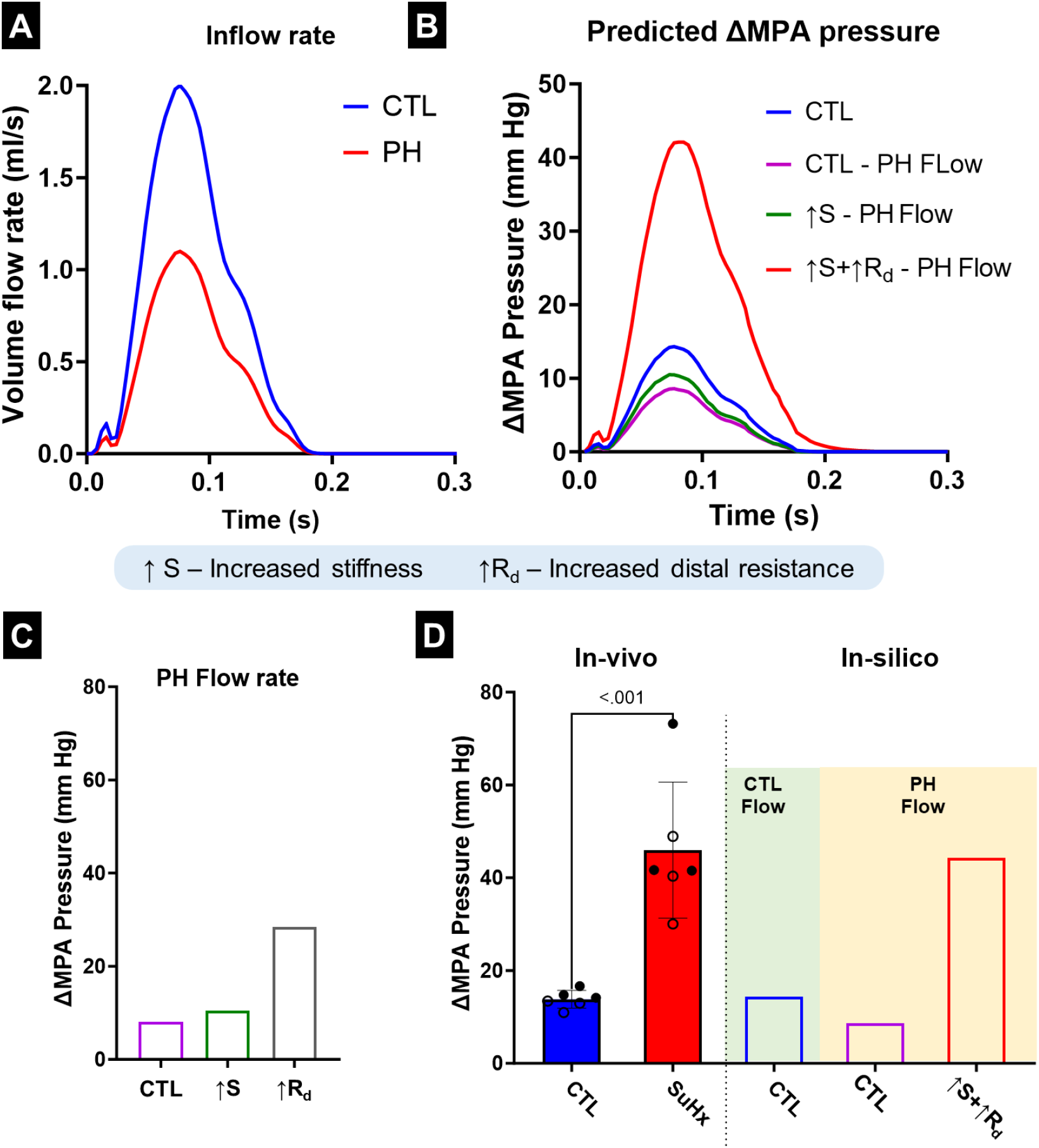
**(A)** Measured main pulmonary artery (MPA) flow profile in the control (CTL) and pulmonary hypertension (PH) rats used in the in-silico model. **(B)** The simulated pressure waveform obtained from the in-silico model for the MPA after altering flow and remodeling parameters. **(C)** Simulated changes in ΔMPA pressure as a result of individual remodeling parameters. **(D)** The ΔMPA pressure obtained through in-vivo measurements and in-silico simulations. Male and female animals are denoted by filled and hollow markers, respectively. CTL: n=6, PH: n=6. ΔMPA Pressure: difference in the MPA pressure between systole and diastole.

When investigating the downstream pressure drop in the complete vasculature, a sharp decrease was observed in the pressure after 3-4 generations in both the healthy and PH geometries (Figs. 6A, B). The flow rate in the vasculature split at the branching points in the same ratio as the lumen area of the daughter vessels (Figs. 6C, D). This led to the two primary branches carrying the majority of the flow, and there was a significant decrease when the flow reached the smaller vessels.

**Fig. 6.**
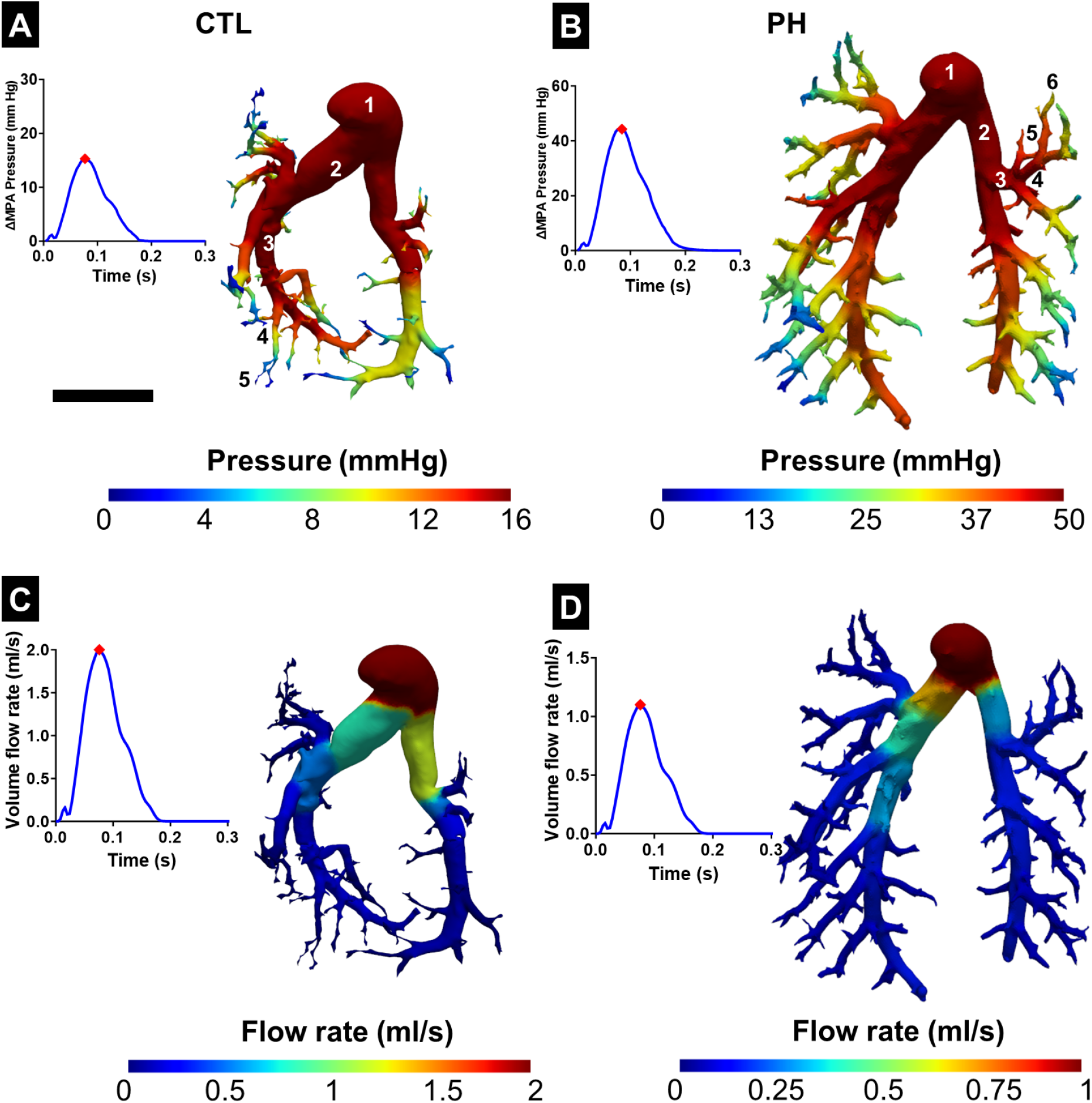
Variation of pressure in the pulmonary vasculature in **(A)** control (CTL) and **(B)** pulmonary hypertension (PH) rats with the corresponding time point marked on the pressure-time plot. Variation of flow rate in the pulmonary vasculature in **(C)** CTL and **(D)** PH rats. Results from the 1D FSI simulation were mapped onto the 3D geometry obtained from segmentation with the corresponding time point marked on the flow-time plot. Representative branching generations of the vessel have been labeled in (**A**) and (**B**). ΔMPA Pressure: difference in the MPA pressure between systole and diastole. Scale bar represents 5mm.

### 3.4 Effect of PA remodeling on PA impedance

*Z*_0_ and *Z*_*c*_ were estimated from the variation of PAZ with frequency (Fig. S3). Frequency domain analysis indicated that increased resistance was the dominant contributor towards *Z*_0_ (Fig. 7A), with *Z*_0_ due to increased resistance being 82.7% of the *Z*_0_ in PH. In contrast, increased stiffness had a larger contribution towards *Z*_*c*_ compared to increased resistance (Fig. 7B), with *Z*_*c*_ due to increased resistance being 41% of the *Z*_*c*_ in PH. The impedance frequency analysis also indicated an interaction between increased stiffness and resistance, with the *Z*_0_ and *Z*_*c*_ values estimated from the in-silico PH simulation not being the arithmetic sum of the impedance values estimated in the cases of the individual remodeling events. In addition, *Z*_0_ and *Z*_*c*_ estimated through in-silico modeling closely matched the mean *Z*_0_ and *Z*_*c*_ values estimated through catheterization (Figs. 7A, B).

**Fig. 7.**
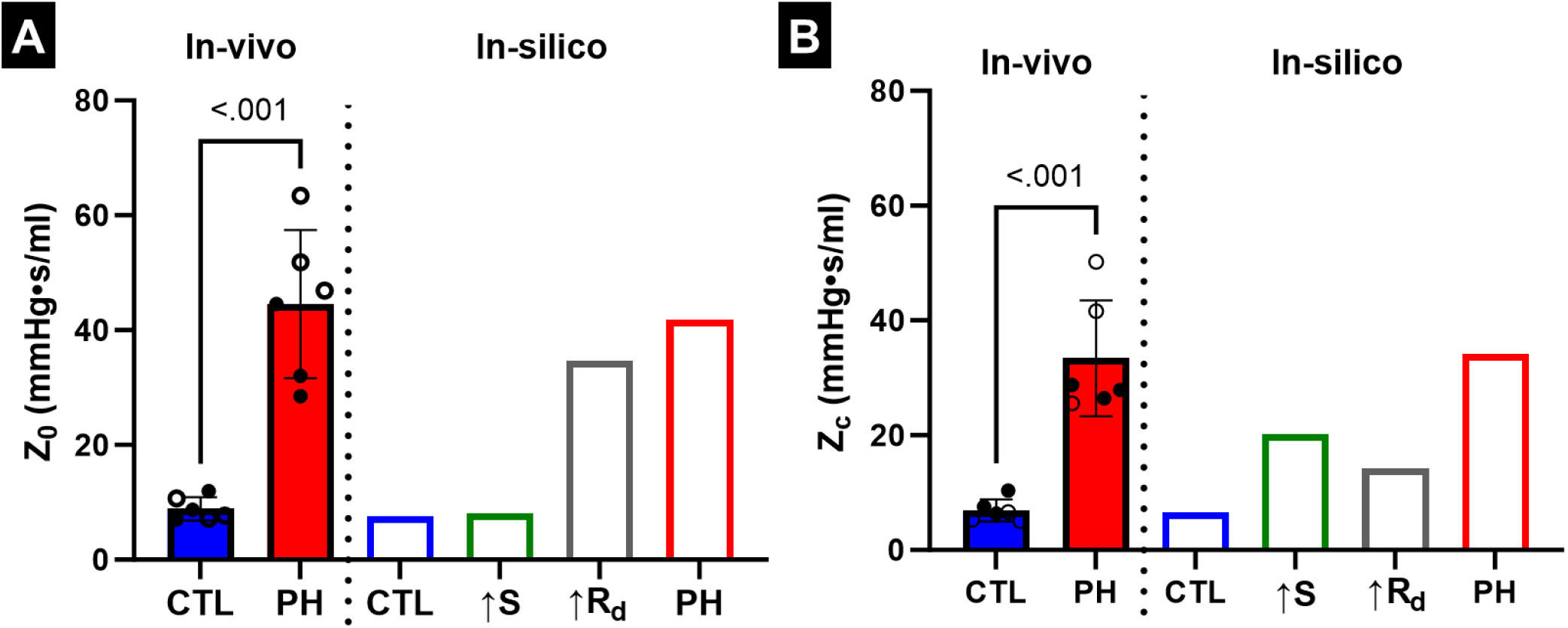
**(A)** Variation of 0Hz PA impedance (*Z*_0_) due to PA remodeling events. **(B)**Variation of characteristic PA impedance (*Z*_*c*_) due to PA remodeling events. Male and female animals are denoted by filled and hollow markers, respectively. CTL: n=6, PH: n=6.

### 3.5 Validation of 1D FSI model and effect of non-linear vessel behavior

The model was validated by simulating a constant volume flow rate through a near-rigid tube (detailed in Section S.5 in the supplementary material). The Windkessel resistance values were set to 0 (i.e., zero outflow pressure). We observed that the pressure contour estimated by the 1D FSI model matched with the result provided by a commercial computational fluid dynamics software (3D-FSI using ANSYS Fluent and Structural; Figs. S4A, B). We also compared the effect of linear and non-linear vessel behavior as a function of the expansion of the lumen area. We observe that the behavior is similar under small values of area deformation, but the non-linear vessel behavior presented in section 2.4 leads to significantly larger pressure values at large values of lumen area deformation (Fig. S5). This indicates a need for a significant increase in RV afterload to maintain similar flow behavior in the pulmonary vasculature.

## 4 Discussion

### 4.1 Deconvoluting the effects of vascular remodeling events on MPA pressure

We have presented a method to investigate the subject-specific contribution of increased resistance and vessel stiffness to the increased MPA pressure in a PH model of rats. The results of this study indicate that the remodeling mechanisms have complex interplay and that summing the individual effects of each mechanism fails to fully capture the final PH state. This highlights the utility of the model presented in this study in investigating and estimating the effect of each mechanism, as well as their interactions with each other. Under a fixed input flow rate at the MPA, each individual vascular remodeling event resulted in an increase in ΔMPA pressure. This indicates that increased distal resistance and increased stiffness in both the proximal and distal vessels each contribute to increased MPA pressure. Increased resistance had the greatest effect on increasing the ΔMPA pressure. This is consistent with the use of PVR as a key longitudinal metric of PH. Increased proximal and distal stiffness affected pressure similarly; their effect on pressure was smaller than that of resistance.

As previously stated, the final PH ΔMPA pressure was not the result of the sum of the changes from each individual mechanism, suggesting a significant interaction taking place between mechanisms. Interactions between remodeling events were most evident in the relationships between proximal stiffness and other mechanisms. Increased proximal stiffness had greater effects on peak ΔMPA pressure when present in conjunction with increased resistance. Indeed, these results highlight the complex nature of post-PH remodeling and point towards a need for the continued exploration of patient-specific modeling approaches to identify the most effective therapeutic plan in each case.

### 4.2 Effect of vascular remodeling on pulsatility of flow

Notable changes were observed in the pressure waveform due to remodeling events, given a fixed volume flow rate waveform. These variations in the pressure waveform were reflected in the 0Hz (*Z*_0_) and characteristic (*Z*_*c*_) PA impedance values estimated through Fourier analysis. Studies have indicated that PA impedance is a more comprehensive biomarker than PVR due to its ability to capture both static and pulsatile components of the pressure and flow waveforms^39, 41^. The study by Hunter et al.^41^ indicated that *Z*_0_ demonstrated a significant correlation with PVR, while *Z*_*c*_ correlated with decreased vessel compliance. The results of our current study are in agreement with these studies, with increased resistance being the dominant contributor towards increased *Z*_0_, and increased stiffness being the major contributor towards increased *Z*_*c*_. Also, studies have reported that PA impedance measurements correlate better with patient outcomes in PH^39, 41–43^, corroborating that peak pressure is insufficient to understand the degree of remodeling and its effect on pulmonary circulation.

In addition, there were several qualitative variations in the pressure waveform due to the vascular remodeling events. In the case of increased flow resistance, there was a delay between the peaks in volume flow rate and pressure. In addition, increased resistance also resulted in a significantly slower pressure drop, with pressure requiring the duration of one volume flow pulse to drop to EDP. These variations are hypothesized to be a result of a combination of pruning (reduction in the number of arterioles and alveolar capillaries)^1, 4^ and increased resistance in the distal vessels impeding fluid flow, leading to the accumulation of fluid. This accumulated fluid slowly exits the vessel, thus leading to an “uncoupled” behavior with the volume flow rate waveform. However, incorporating increased vessel stiffness in addition to the increased resistance reduced the effect of both the delay between pressure and volume flow rate waveforms and a slower rate of pressure drop. This was expected as the increased vessel stiffness prevented excessive vessel dilation, effectively forcing the fluid downstream.

### 4.3 Capturing PA remodeling using a non-linear and anisotropic material model

In this work, the H-O (non-linear anisotropic) constitutive model was used to describe the mechanical behavior of the arterial vessels. To our knowledge, this is the first example of the incorporation of a non-linear anisotropic material model that is expressly designed to model arterial behavior into a 1D FSI arterial modeling approach. While vessel behavior has been shown to be non-linear, linear constitutive models are commonly used for their simplicity. The primary difference between linear and non-linear vessel behavior is the capability of non-linear behavior to capture the significantly increased pressure at large values of lumen area expansion. Vascular remodeling events, such as narrowing of the distal vasculature, arterial hypertrophy, and pruning, lead to the reduction of the lumen cross-sectional area, leading to a larger area expansion under similar vascular blood flow. The predicted pressure as a result of larger area deformation of the vessels is not capable of being accurately captured by a linear behavior regime and is more accurately represented by a non-linear vessel model. In addition, the increased arterial wall stiffness leads to increased stiffness of the arteries and arterioles. This increased stiffness, combined with the decreased lumen area, will lead to an increased flow rate through the arterioles and capillaries, reducing oxygen diffusion into and carbon dioxide out of the bloodstream. This effect manifests as shortness of breath, which has been reported to be a symptom of PH by several studies^17, 44, 45^.

In addition to non-linear mechanical behavior, the incorporation of material anisotropy is also essential when investigating the vascular remodeling observed in PH. Our results indicated that the stiffening of the vessel walls is not uniform in the circumferential and longitudinal directions. The MPA from healthy rats was mildly biased towards the circumferential direction, and the MPA from the PH rats was further biased towards the longitudinal direction. These results indicate fiber re-orientation in the vessels due to PH and emphasize the need for an anisotropic material model. The alteration in vessel wall anisotropy also highlights a potential opportunity to introduce longitudinal dependence on pressure in the solid domain of the 1D FSI model.

### 4.4 Implications for potential clinical application

A combination of in-vivo and ex-vivo measurements that characterize the increase in MPA pressure and vascular impedance due to PH were incorporated into an in-silico model. Then, *hypothetical states* were generated in-silico to analyze the isolated effect of individual remodeling mechanisms on ΔMPA pressure. This modeling approach can enable an improved subject-specific understanding of the interaction between vascular remodeling events and between individual events on RV and PA sides^46, 47^, and serve as an essential step toward the optimization of clinical intervention strategies. However, the data acquisition methods used in this study to obtain experimental measurements would require modifications for application in the clinic. In the case of human patients, the vessel stiffness and overall vascular resistance can be estimated through the measurement of PAC and PVR using echocardiography and catheterization, respectively^48, 49^. The structure of the pulmonary vasculature can be reconstructed through pulmonary angiography. Generation of the hypothetical states presented in this study can then allow for individualized risk stratification, leading to improved clinical therapeutic strategies.

### 4.5 Limitations

In this study, the radius measured through *µ*CT imaging was assumed to be the same as the radius in the load-free configuration. However, this radius may not accurately reflect the radius of the vessels in vivo due to two factors. Firstly, the fixation is performed at elevated pressure (15 mmHg)^34^ to allow for the imaging of smaller vessels, but this can lead to the expansion of vessels and the overestimation of radii. Secondly, the measured radius from *µ*CT may be susceptible to errors from the resin filling, imaging, and post-processing. This factor will primarily affect smaller vessels, which may not be fully infused with resin, and the radius may be incorrect due to imaging resolution and processing artifacts. A limitation of the constitutive model and 1D FSI framework used in this study is that pressure is dependent on only the circumferential and radial stresses. However, the fibers in the vessel wall are also partially oriented along the axial directions. This implies that pressure at a given cross-section would also be a function of the pressure in the axial surroundings. The lack of this constraint can allow for sudden changes in the lumen area along the axial length of the vessel. This constraint will be incorporated in future studies to ensure physiologically reasonable deformation of the artery along the axial direction in addition to physiologically reasonable deformation of the lumen area.

## 5 Conclusions

In this study, we presented a computational approach to separate the effects of multiple vascular remodeling events in the pulmonary arteries towards the altered MPA pressure amplitude and pulsatility associated with PH. We combined in-silico modeling with in-vivo and ex-vivo measurements from healthy and diseases rats to generate hypothetical states in-silico of the increase in MPA pressure caused by a single factor. The results of our study indicated that the increased vascular resistance is the dominant contributor towards increased pressure amplitude, while increased vessel stiffness is the major contributor towards altered pulsatility. The method presented in this study can improve our understanding of the effect of vascular remodeling events on altered pulmonary hemodynamics, and serve as an essential step towards the individualization and optimization of clinical intervention strategies.

## 6 Declaration of competing interest

The authors declare no conflict of interest.

### 7 Acknowledgements

This work was supported by the National Institutes of Health (R00HL138288 to R.A.)

## 8 Data availability

The data that support the findings of this study are available from the corresponding author, R.A., upon request.

## Supplementary Material

### S.1 Non-linear and anisotropic constitutive model for the pulmonary artery

The isotropic and anisotropic strain energy functions are given by:

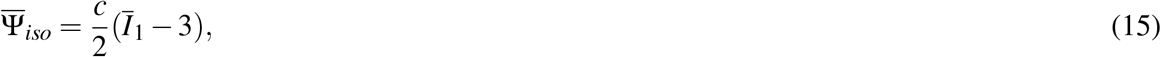

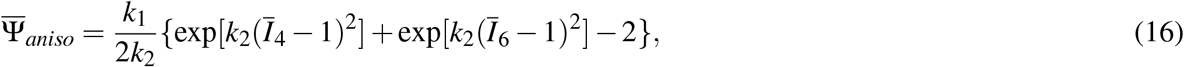

where *c, k*_1_ and *k*_2_ are material parameters. *c* is associated with the isotropic (ECM) response, while *k*_1_ and *k*_2_ are associated with the fiber (anisotropic) response. The Cauchy stress is given by

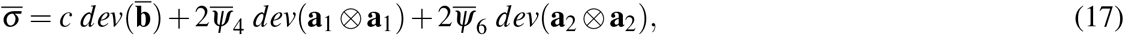

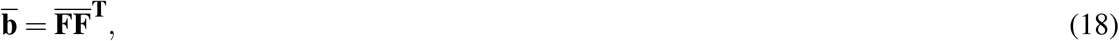

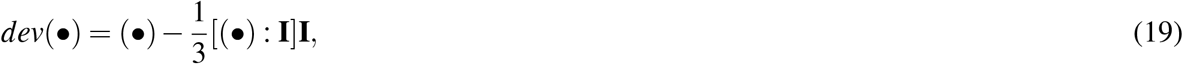

where 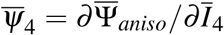 and 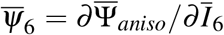 are the derivatives of the anisotropic strain energy function with respect to the invariants 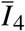 and 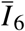. 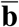 is the left Cauchy-green tensor. The derivatives were computed to be

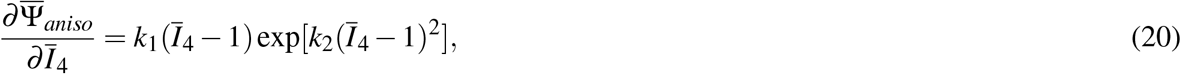

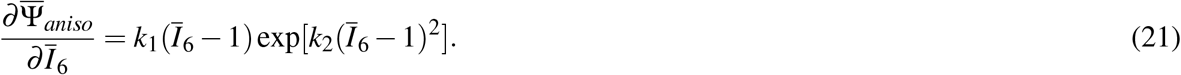

For an ideal case, assume the following isochoric deformation vector with no shear components

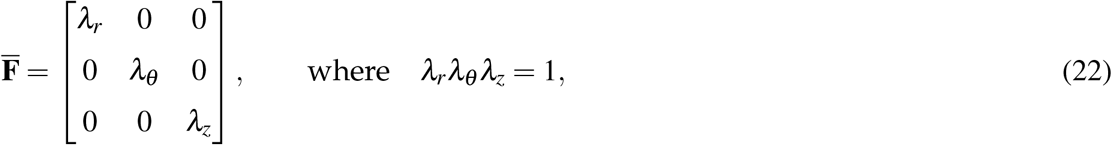

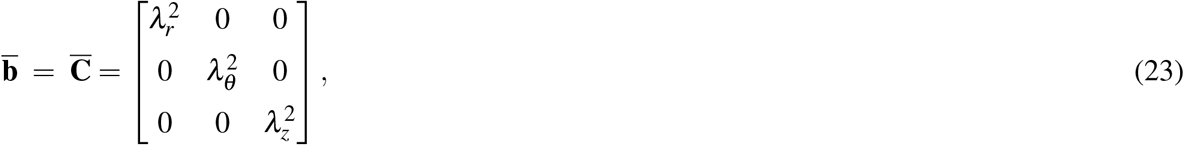

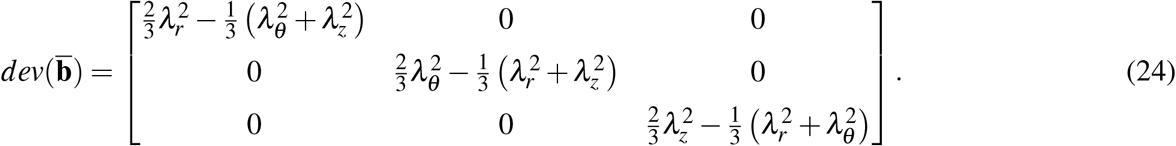

Hence, the isotropic stress is given by

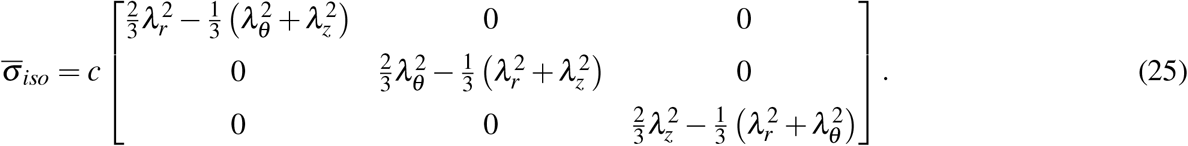

Considering the effects of the fibers, 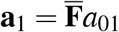, where *a*_01_ and *a*_02_ are given by the fiber angle *β*.

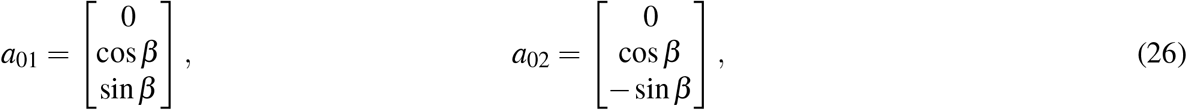

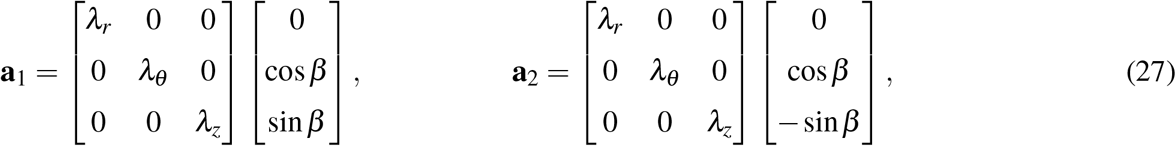

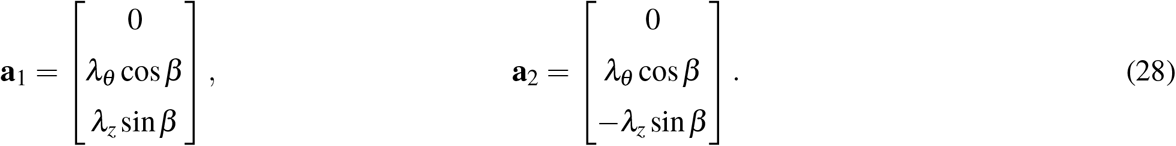

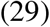

Now converting **a**_1_ and **a**_2_ into matrices and determining the deviatoric part

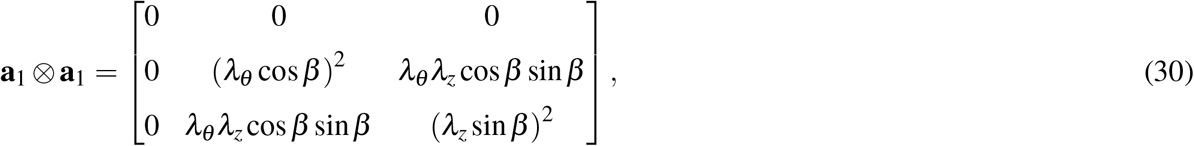

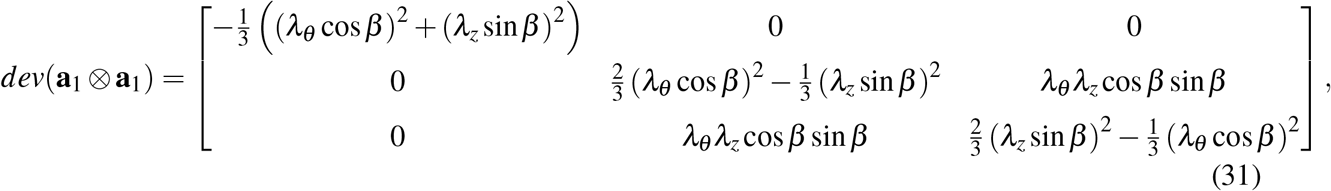

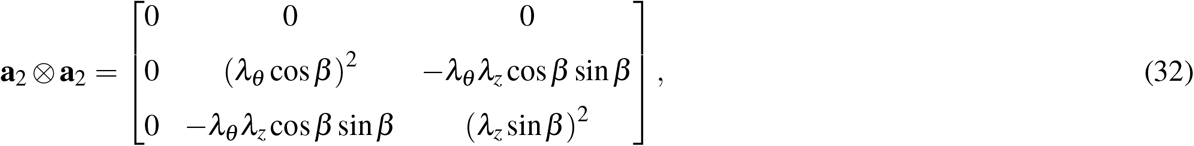

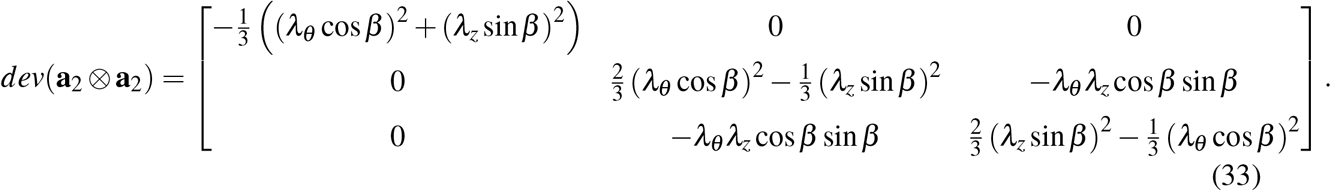

Now computing the stress *σ*_*θθ*_

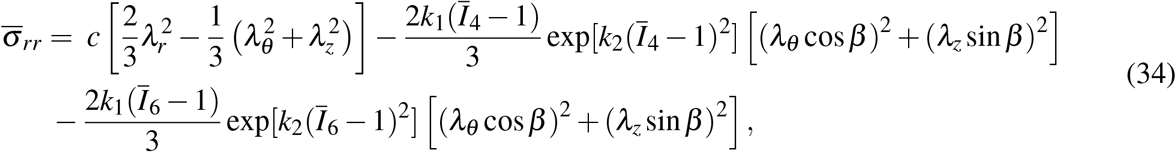

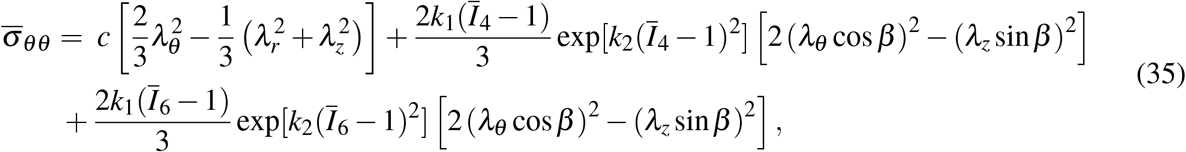

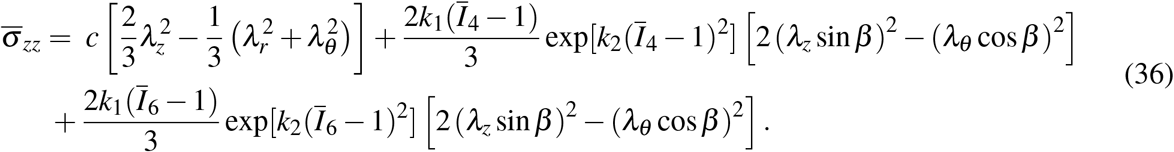

Now computing the fiber-related invariants

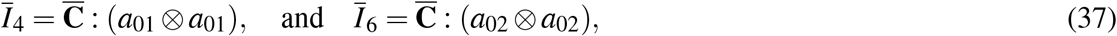

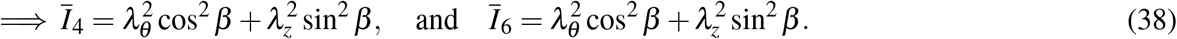

The final stress values are

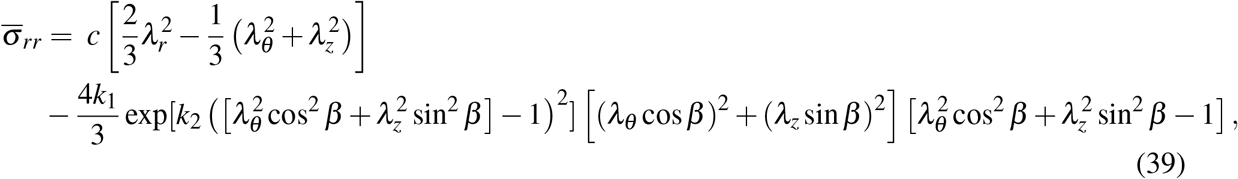

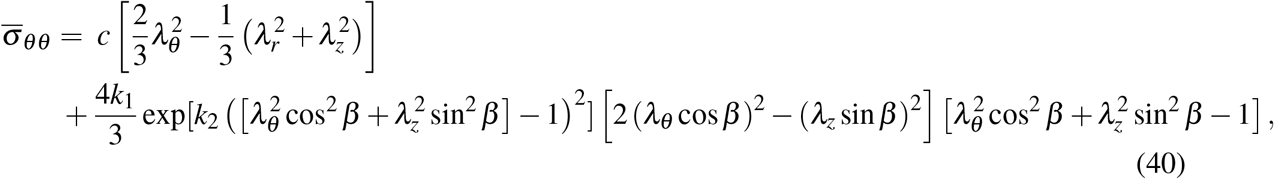

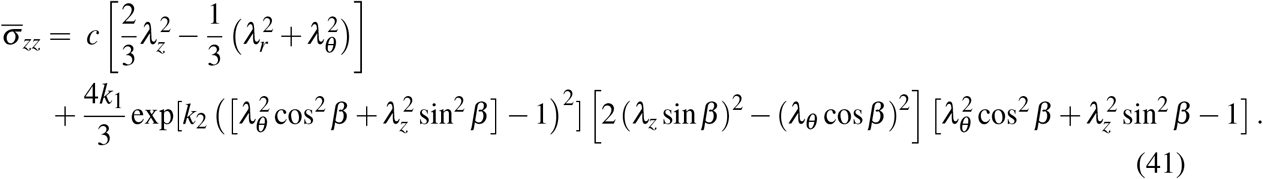

We know that 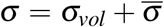, where *σ*_*vol*_ = *p***I** is the hydrostatic stress and *p* is the hydrostatic pressure.

### S.2 Fluid pressure during inflation of vessel

For pure inflation of a cylinder, the variables in the deformed configuration are given by *r* = *r*(*R*), *θ* = Θ, and *z* = *Z*. Here *R ∈* [*R*_*i*_, *R*_*o*_], Θ *∈* [0, 2*π*] and let *R*_*o*_ − *R*_*i*_ = *h* be the thickness. Hence, the deformation gradient and Cauchy green tensors are given by

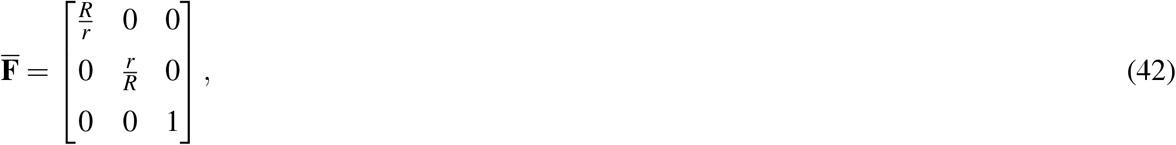

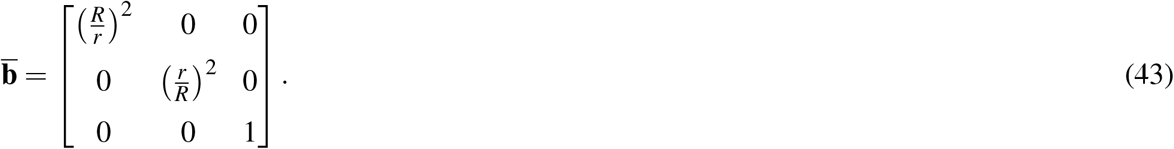

We can compute hydrostatic pressure using the equilibrium equation ∇*σ* = 0. In cylindrical coordinates

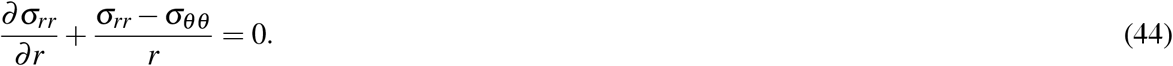

We use this to get the value of internal pressure as

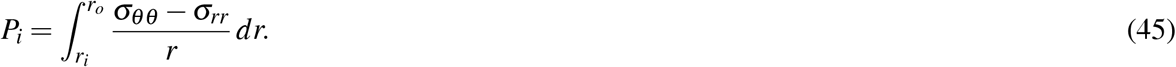

We can substitute 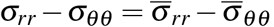, because 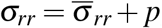, and 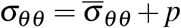, where *p* is the hydrostatic pressure. Hence

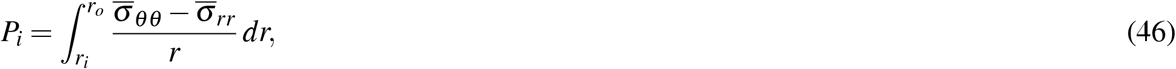

Computing the RHS

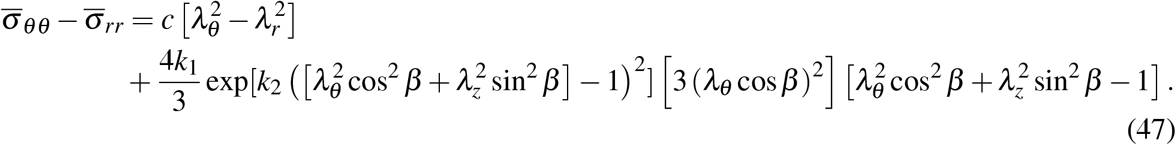

Substituting the values for *λ*_*r*_, *λ*_*θ*_, *λ*_*z*_, we get

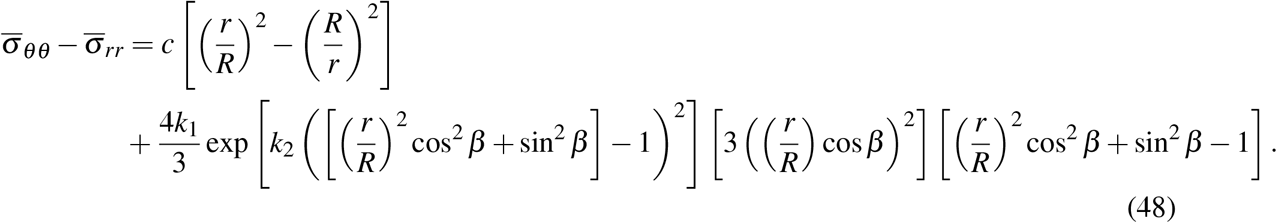

We can compute *P*_*i*_ by integrating Eq. S46 over the thickness of the vessel.

### S.3 Uniaxial testing behavior

**Figure S1.**
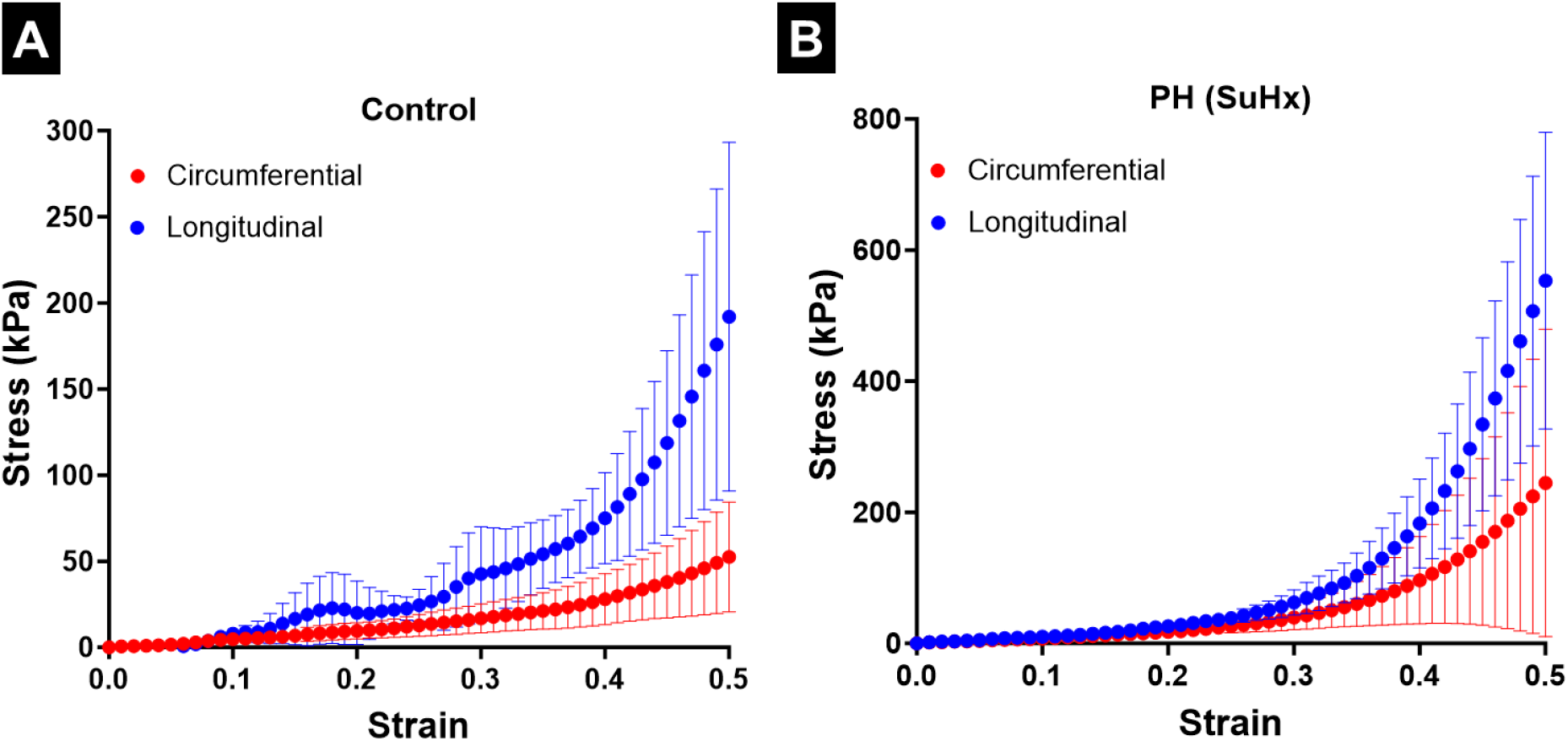
Uniaxial testing of MPA tissue specimens from (**A**) control rats, (**B**) PH rats. CTL: n=6, PH: n=6.

### S.4 Effect of remodeling parameters on pressure waveform

**Figure S2.**
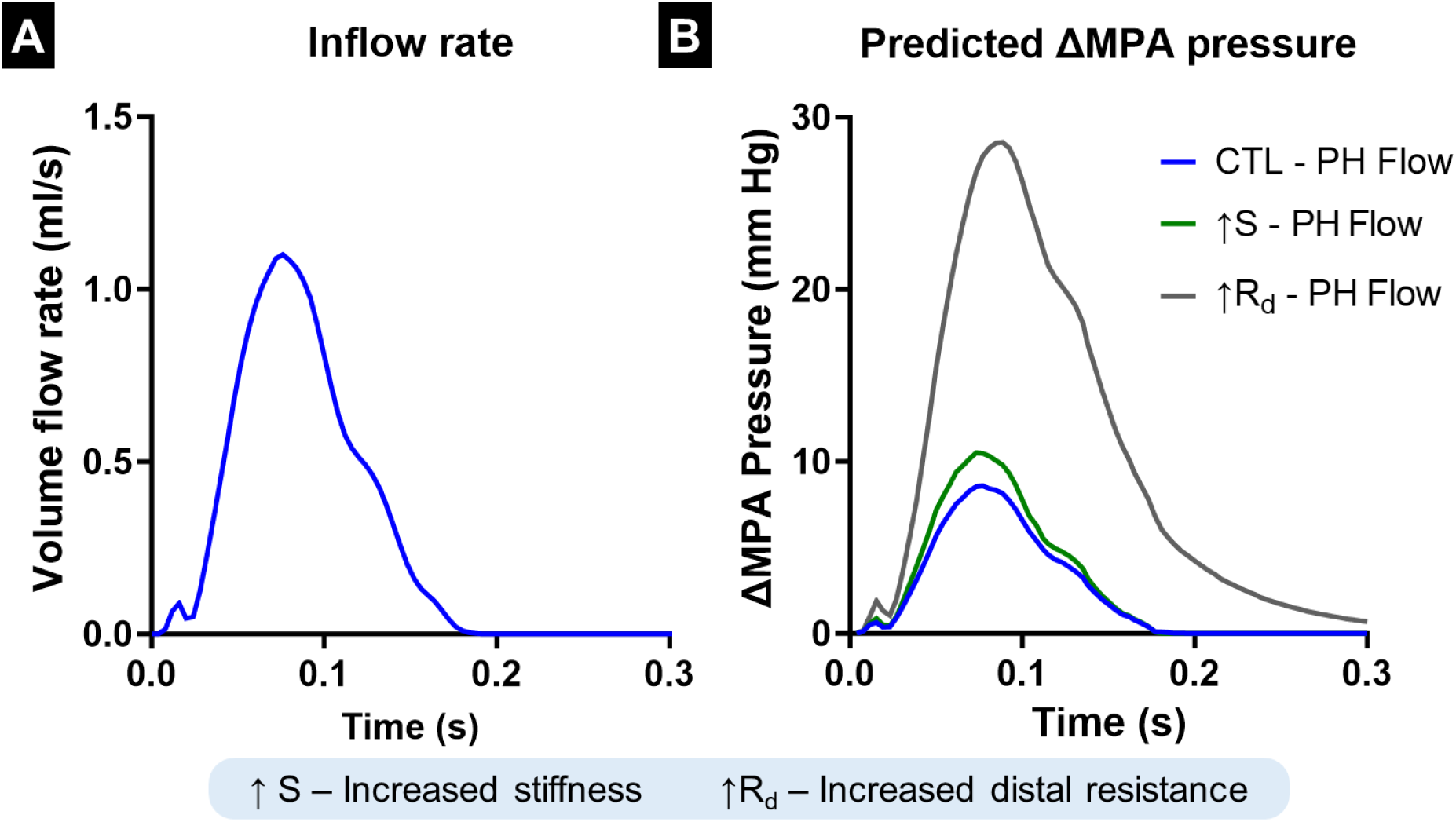
Effect of individual vascular remodeling events on the pressure waveform. (**A**) Volume flow rate at the inlet, (**B**) Pressure waveform for increased vessel stiffness and distal resistance. (**C**) Volume flow rate at the inlet with an increased period of zero flow after the pulse, (**B**) Pressure waveform increased distal resistance to study the relaxation behavior.

### S.5 Variation of PA impedance with frequency

**Figure S3.**
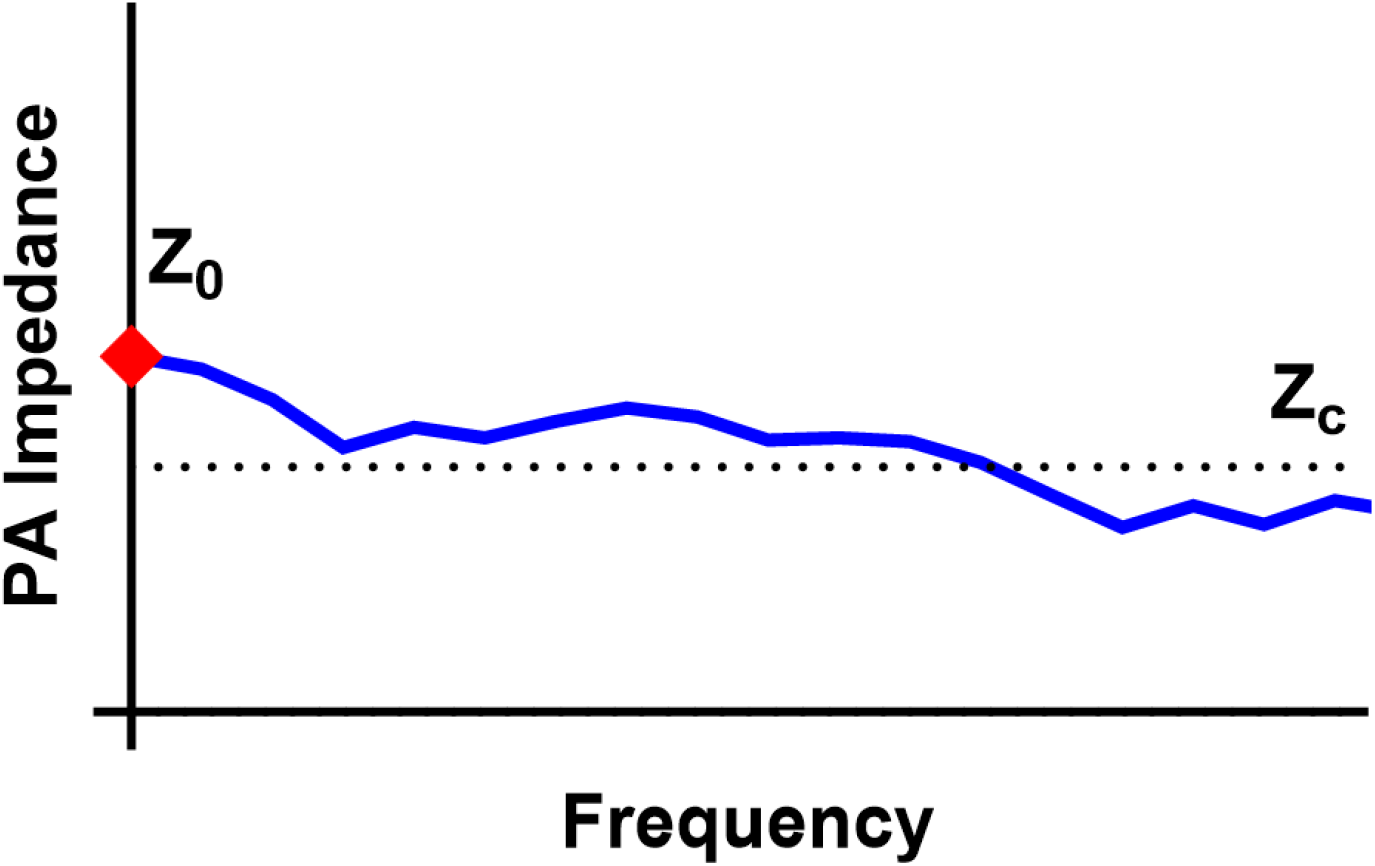
Schematic describing the variation of impedance with frequency and the estimation of 0Hz-and characteristic impedance.

### S.6 Validation against 3D FSI

For validation, we have compared the 1D FSI framework presented in this study against a 3D FSI framework. Here we simulated a constant fluid flow rate through a Y-branch, where the length and radius of each vessel are 50mm and 5mm respectively. The thickness is set to be 20% of the radius (1mm). The flow rate is set to 30ml/s. For the vessel properties, the stiffness of fiber components was set to 0, and the shear modulus (*c*) was set to 60kPa. To perform the equivalent simulation in 3D, we performed the simulation in ANSYS workbench, coupling the transient structural and Fluent modules. Here, the displacement of the inlet and outlet faces was constrained such that only radial and circumferential displacement was allowed. A single additional point on the inlet was completely fixed to ensure stable simulations.

**Figure S4.**
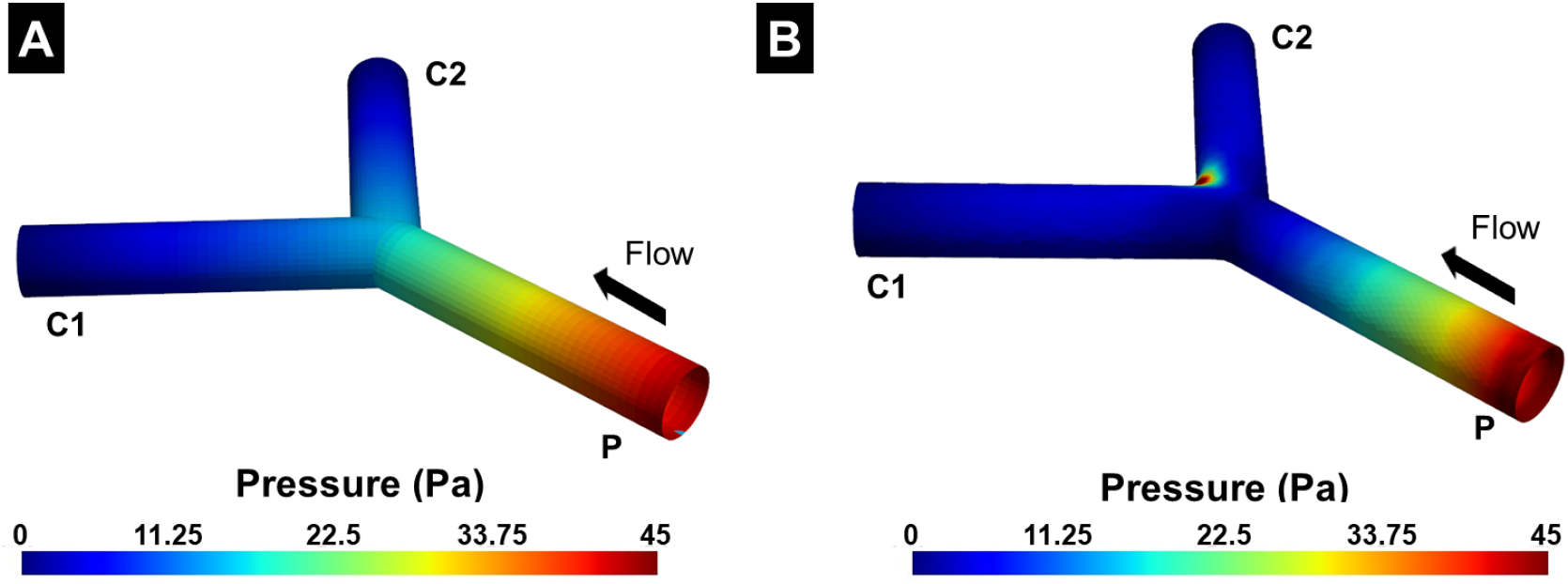
(A) The pressure predicted by the 1D simulation mapped onto a representative geometry, and (B) Pressure on the wall of the fluid domain. **P**: Parent vessel; **C1, C2**: child vessels.

The results indicated good agreement between the 1D and 3D simulations. The pressure at the inlet in the 1D simulation was 42.8 Pa. Correspondingly, the pressure on the fluid domain wall from the 3D simulation was 44.7Pa. The difference between the predicted values is likely due to the idealized bifurcation assumed by the 1D FSI framework.

### S.7 Effect of non-linear material model

**Figure S5.**
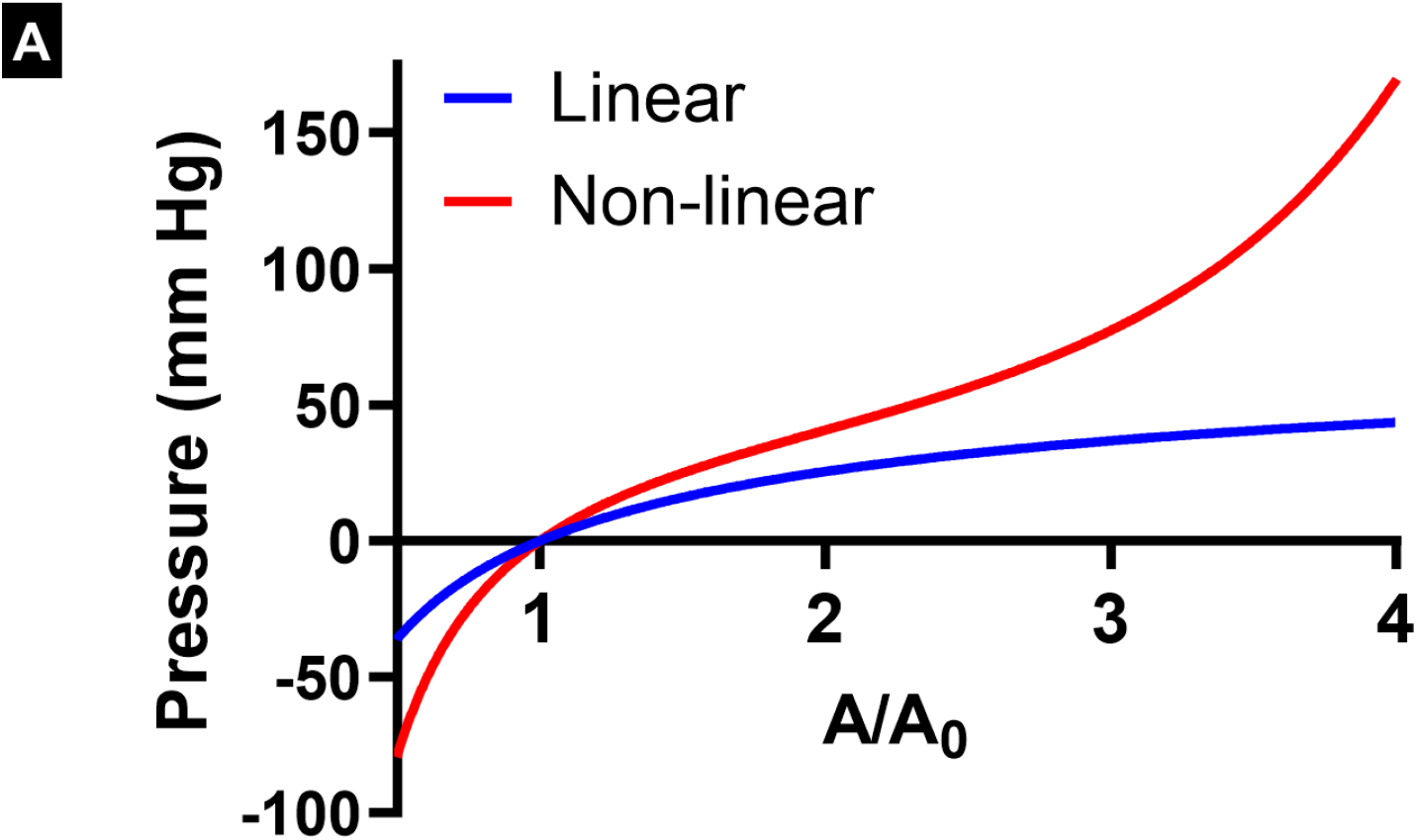
Variation of pressure with lumen area. Lumen area was normalized by the original lumen area

## Notes

### Competing Interest Statement

The authors have declared no competing interest.

